# Sex Differences in Contextual Extinction Learning After Single Binge-Like EtOH Exposure in Adolescent C57BL/6J Mice

**DOI:** 10.1101/2024.10.25.620195

**Authors:** Kiara M. Cardona-Jordan, Xiany X. Lay-Rivera, Eliezer Cartagena-López, Dina L. Bracho-Rincón, Ruth González-Bermejo, Gerardo L. Alvarado-Monefeldt, Jovangelis P. Gonzalez Del Toro, Christian J. Esquilín-Rodríguez, Mario Lloret-Torres, Cristina Velázquez-Marrero

## Abstract

The relationship between chronic heavy drinking and post-traumatic stress disorder (PTSD) is well-documented; however, the impact of more common drinking patterns, such as a single episode leading to a blood alcohol concentration (BAC) of 0.09 g/dL (moderate intoxication), remains underexplored. Given the frequent co-occurrence of PTSD and alcohol misuse, it is essential to understand the biological and behavioral factors driving this comorbidity. We hypothesize that alcohol’s immediate sedative effects are coupled with the development of persistent molecular alcohol tolerance, which may disrupt fear extinction learning. To investigate this, we employed a ***S****ingle **E**pisode **E**thanol* (SEE) *in-vivo* exposure to mimic binge-like alcohol consumption over a 6-hour period, following contextual conditioning trials. Extinction trials were conducted 24 hours later to assess the effects on extinction learning. Our findings reveal a significant deficit in fear extinction learning in alcohol-treated adolescent male mice compared to saline-treated controls, with no such effects observed in female adolescent mice. These results suggest that even non-chronic alcohol exposure may contribute to the development of trauma- and stress-related disorders, such as PTSD, in males. Additionally, histological analysis revealed significant alterations in FKBP5, β-catenin, and GSK-3β levels in the hippocampus, striatum, and basolateral amygdala of alcohol-treated mice following extinction. The insights gained from this study could reshape our understanding of the risk factors for PTSD and open new avenues for prevention and treatment, targeting the molecular mechanisms that mediate alcohol tolerance.

**STATEMENT OF SIGNIFICANCE:** This study investigates the impact of binge-like alcohol exposure on context extinction learning, aiming to identify previously unrecognized risks associated with this common drinking pattern and the development of trauma- and stress-related disorders, such as PTSD. Our findings reveal that binge-like alcohol exposure impairs extinction learning in male adolescent mice by disrupting molecular mechanisms within fear memory circuits, suggesting novel therapeutic and preventive targets. Dysregulated candidates include the canonical Wnt/β-catenin signaling proteins, β-catenin and GSK-3β, along with FKBP5, a key player in glucocorticoid signaling and part of a gene network linked to PTSD. These alterations, found in the dorsal hippocampus (dHPC), basolateral amygdala (BLA), striatum, and nucleus accumbens (NAc) core and shell, may serve as promising targets for future pharmacological intervention.

## INTRODUCTION

Adolescence is a critical developmental stage during which exposure to stressful events can create risk factors for psychiatric disorders, including alcohol use disorders and anxiety. Binge drinking is the most common pattern of alcohol consumption among adolescents and young adults (1–5). The National Institute of Alcohol Abuse and Alcoholism (NIAAA) defines binge alcohol consumption as an episodic pattern of drinking that brings blood alcohol concentration (BAC) to 0.08 percent or higher (1, 3, 6–9). For males, this corresponds to consuming five or more alcoholic drinks in a single episode; for females, it is four or more drinks. Research indicates that repeated episodes of binge drinking during adolescence can alter adolescent brain development and cause severe deficits in social, attention, memory, and other cognitive functions (10–15). During this critical period of brain maturation, cognitive, emotional, and social development can be significantly impacted by ethanol exposure (10, 13, 15–20). Numerous studies have shown a connection between early life stress and increased rates of alcohol use disorders (AUDs) and post-traumatic stress disorder (PTSD) (16, 17, 21–24). Although adolescents and young adults who engage in binge drinking represent a key demographic for research, the effects of a single binge episode on fear extinction learning remain largely unexplored.

Binge drinking in adolescents impacts critical brain structures, including the hippocampus, prefrontal cortex, frontal lobe, and amygdala (25). These regions are vital for both long-term and short-term memory and their consolidation. Additionally, chronic ethanol consumption is associated with impairments in memory, learning, impulse control, and the balance between rational decision-making and emotional responses, as well as trauma- and stressor-related disorders like PTSD (26, 27). PTSD has a high comorbidity with alcohol use disorder (AUD) and is a mental health condition characterized by the brain’s re-experiencing of traumatic events, leading to intense feelings of fear, anger, stress, and anxiety mainly triggered by contextual cues (28–31). Our study emphasizes the established connection between the persistence of traumatic memories and deficits in extinction learning. Clinical research has indicated that alcohol use disorder is associated with fear memories, demonstrating that extinction learning is slower, weaker, and less context-specific under the influence of alcohol, resulting in a more enduring fear response (32). This underscores the importance of understanding the role and potential mechanisms by which binge alcohol exposure affects context extinction learning in both male and female mice.

Wnt/β-catenin activation reduces the surface expression of large-conductance calcium- and voltage-activated potassium (BK) channels, which correlates with increased β-catenin expression in both the hippocampus and striatum. This alteration likely impacts the necessary changes in intrinsic excitability required for contextual extinction learning (33–36). Consequently, we investigated β-catenin protein expression following 24 hours of extinction trials. We also examined the expression of FKBP5, a gene that encodes a molecular co-chaperone of the glucocorticoid receptor, which modulates intracellular glucocorticoid signaling and plays a crucial role in regulating the stress response (37–41). FKBP5 influences the sensitivity of glucocorticoid receptors (GR) through its interactions with co-chaperones of the steroid receptor complex, leading to a prolonged stress response (37, 38, 40). The FKBP5 gene, which encodes FK506-binding protein 5, is implicated in various trauma- and stress-related disorders, as well as alcohol use and withdrawal (42, 43). Additionally, FKBP5 gene expression levels have been associated with PTSD (41, 44) and the regulation of fear memories (39, 45). Furthermore, longitudinal studies examining stressors and alcohol exposure during early development have shown increased expression levels of this gene (46), suggesting a potential link between early alcohol and/or stress exposure and future epigenetic changes. Targeting the dysregulation of these genes has emerged as a potential clinical strategy for preventing PTSD (47, 48) which now may apply to prevention of comorbid diseases linking *non-chronic* common alcohol use patterns and trauma responses in adolescents.

## METHODS

### Subjects

We used C57BL/6J adolescent male and female mice (P30-P45) (Jackson Laboratories, MO), 120-140 mice were individually housed in ventilated cages fitted with steel lids, filter tops, and with unrestricted access to food and water on a 12/12-hour light/dark cycle. At all times, animals were treated in accordance with the National Institutes of Health, Guide for the Care and Use of Laboratory Animals (Institute of Animal Resources, 1996). Mice were handled and housed in accordance with the standards established by the Institutional Animal Care and Use Committee of the University of Puerto Rico Medical Sciences Campus and the National Institutes of Health. All experimental procedures were approved by the Institutional Animal Care and Use Committee (IACUC).

### Single Episode Ethanol (SEE) exposure

Mice were equally distributed by weight and freezing behavior learning during the Contextual Fear Conditioning phase, between the saline (control group) and ethanol group. One of the main limitations in the study of binge drinking is that animal models will not voluntarily drink the necessary amount to reach the required blood alcohol concentrations (49). To avoid this issue mice received six intraperitoneal injections (i.p.) of 20% EtOH v/v with saline solution using a 26.5-gauge needle evenly spaced over 6 hours, after contextual fear conditioning trials. The first injection was at an EtOH concentration of 1.8 g/kg, and the subsequent five injections were at a concentration of 1.2 g/kg. After each injection, mice were placed back in their housing and carefully monitored. Blood alcohol concentrations returned to naïve levels after 3hr withdrawal as previously described in (50).

### Contextual Fear Conditioning and Extinction Paradigm

Contextual fear conditioning and extinction were performed during the animal’s light cycle in a training cage within a sound attenuating chamber minimizing travel of ultrasonic vocalizations and fear-related pheromones (Stoelting Co, IL). Home cage conditions were carefully chosen and carried out consistently to minimize the negative impact on context fear conditioning learning as in (51). Behavior, tracking, and freezing of the animal were recorded by low-light video cameras and ANY-maze software (Stoelting Co, IL). Stimulus (shock) presentation was automated by ANY-maze software. The cages were cleaned with 70% isopropyl alcohol before and after each experiment. Also, mice were tested in the same chamber using the same context in every trial. Before the mice passed through any behavior test, they had a period of 45-60 minutes for acclimation.

#### Contextual Fear Conditioning

Fear conditioning trials were performed in a training cage [17 cm (w) x 17 cm (d) x 25 cm (h)] equipped with stainless-steel shocking grids connected to a precision feedback current-regulated shocker. Protocol followed closely that described in (33). Briefly, mice were placed in Context A and received one 2-second, 0.7mA foot shock 136 seconds into the trial. The trial total time was 228 seconds. After each trial they were returned to their home cages with water and food ad libitum (Fig. 1A).

**Figure 1:**
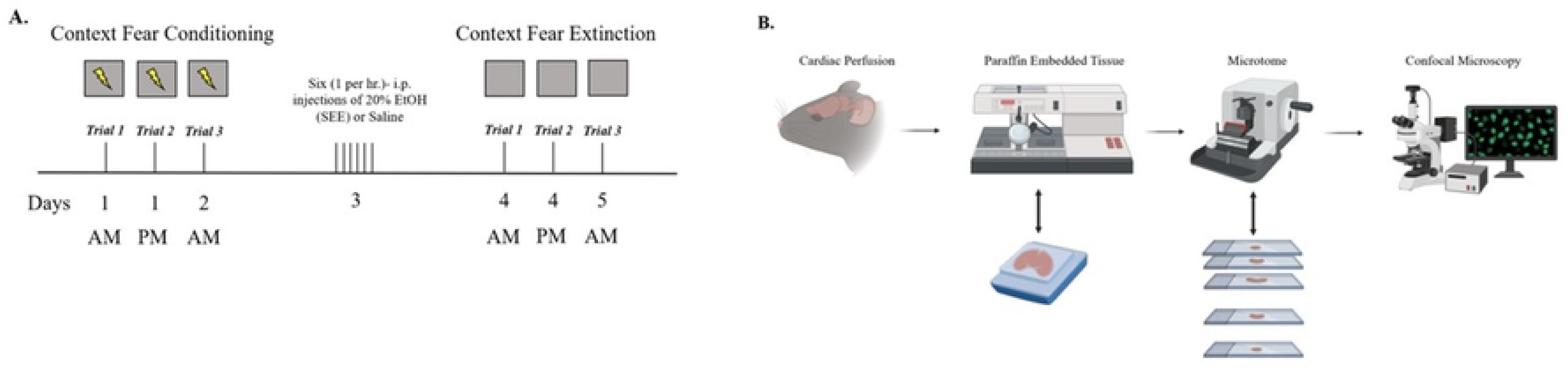
Experimental design. **(A.)** Representative diagrams of the behavioral test in C57BL/6J mice before and after the Single Ethanol Episodic (SEE) exposure. Mice will be tested using contextual fear conditioning and extinction paradigm to examine deficits in fear extinction associated to SEE exposure. **(B.)** After the behavioral experiments, mice will be transcardially perfused to obtain healthy brain tissue. Mice’s brain will be embedded in paraffin, and slices will be created using a rotatory microtome to know the expression of FKBP5, β-catenin, and GSK-3β complex after the behavioral essays. Slices will be stained using primary anti-Fkbp5, anti- β- catenin, anti-GSKβ and with their respective secondary antibodies. Figures created with licensed Biorender.com.

#### Contextual Fear Extinction

Fear extinction trials in the same context were performed in Context A cage without the administration of the shock. For extinction in a novel context, mice were placed in Context B, a cage that differed in appearance from Context A. Mice were in the cages for 3.8 minutes (228 seconds) and then returned to their home cages in both cases. They received three extinction trials (Fig. 1A).

#### Two-Bottle Choice, Every-Other-Day Drinking

Two bottles were prepared for each mouse: Bottle 1: 20% EtOH v/v H₂O solution and Bottle 2: H₂O. Each solution was prepared and separated into 50-mL polystyrene centrifuge tubes topped by rubber stoppers with straight open stainless steel sipper tubes. Both bottles were placed in the cage’s steel lids, filter tops at equal distance and refilled every four days (two “Choice Day”). Bottles alternated sides every choice day. The two bottles were removed a day before every choice day and regular drinking water bottles were provided by the Animal Resource Center of the University of Puerto Rico Medical Sciences Campus and placed in cages. The body weights of mice were recorded every 72 hours, and ethanol and water intake values were recorded every 24 hours (to the nearest 0.2 ml). This data was used to calculate self-administered ethanol dose (i.e., g/kg), water consumption (i,e., mL), and relative preference for ethanol (i.e., 20% ethanol intake/total fluid intake).

#### Open Field Test

Tests were performed in a square chamber (around 40 cm x 40 cm x 40 cm) made from white non-porous plastic. A camera was placed directly overhead of the apparatus for optimal view of exploration. On day 1: habituation, each male adolescent mouse was removed from its home cage and placed in the middle of the empty arena for 5 minutes to allow for free exploration. Training took place on Day 2 (24 hrs after habituation). Training allowed mice free exploration for a minimum of 5 min. On Day 4, the mice were brought from their housing room into the testing room and allowed to acclimate to the procedure room for 45-60 minutes prior to starting the test. A single mouse was removed from the home cage and placed in the middle of the open field maze at a time while concurrently activating the ANY-maze software to begin tracking mouse movement. Free and uninterrupted movement of the mouse throughout the respective quadrant of the maze for a single 10 min period was allowed. At the end of the test period, the mouse was removed from the arena and returned to its home cage. The areas were thoroughly cleaned using 70% v/v isopropyl alcohol. The behavioral assay was repeated with each individual mouse.

#### Paraffin Embedding

Mice were euthanized, and brains fixated using Paraformaldehyde 4% solution in PBS (CAS 30525-89-4) using cardiac perfusion under xylazine/ketamine cocktail anesthesia. After fixation, brain tissue was dehydrated using 50, 75, 95, 100% ethanol solution (Sigma Aldrich CAS 64-17-5) and xylene (Sigma Aldrich 1082984007), immersed in paraffin (Thermo Scientific 22-900-701) at 60°C for 12 hours (paraffin was changed every 4 hours) using the protocol stated in Zhanmu O et al., 2019 (52). After dehydration and paraffin immersion, brain tissue was embedded in paraffin blocks using Epredia™ HistoStar™ Embedding Workstation (Cat. No. A81000002) and were left at 4°C until used (Fig. 1B).

#### Immunohistochemistry

After paraffin embedding brains were sliced using a fully motorized rotary microtome (Leica, RM2250), and slices of the dorsal hippocampus (dHPC) and basolateral amygdala (BLA) brain tissue were collected (5-μm-thick). After all brain slices were made, they were deparaffinized and rehydrated as seen in Zaqout et al., 2020 (53). For permeabilization slides were given 2 washes with PBS 1X (2min), then one wash with PBS 1X (10min), and finally 2 washes with PBS/Gelatin/Triton 0.25% (10 min). Slides then underwent blocking with a 5% BSA solution for 1 hour. After blocking were incubated with primary antibodies for FKBP5 (1:500) (Invitrogen 711292), ß-catenin (1:200) (ab32572), GSK-3ß (1:500) (ab93926) and NeuN (1:500) (ab104224 & ab177487) overnight. The next day slices were washed twice with PBS 1X (10min), then one wash with PBS/gelatin/triton 0.25% (10min). They were then incubated with secondary antibodies, Goat Anti-Mouse Alexa Fluor 488 (ab150113), Goat Anti-Rabbit Alexa Fluor 488 (150077), mounting medium with DAPI (ab104139), Goat Anti-Mouse Alexa Fluor 594 (ab150116) and Goat Anti-Rabbit Alexa Fluor 594 (ab150080). The dilution used for all secondary antibodies was 1:500.

#### Confocal Microscopy Acquisition and Analysis

FKBP5, ß-catenin, and GSK-3ß expressions were measured by a tissue’s fluorescence intensity and were viewed using Nikon Instruments A1 Confocal Laser Microscope using a 10X magnification and analyzed with Nikon Nis Element Advance Research program using the maximum intensity projection (MIP) on all images (Fig. 1B).

#### Statistics

All statistical graphs were generated using GraphPad Prism 7.01 and using repeated measures two-way ANOVA or mixed model followed by a Bonferroni multiple comparison test. The confidence level was set to 0.05 (*p* value), and all results are presented as the mean ± standard error of the mean(54). Significances of these tests are displayed as indicated: * p≤0.05, **p≤0.01, *** p≤0.001 and **** p≤0.0001.

## RESULTS

### Contextual Fear Conditioning

We investigated the effects of binge-like ethanol consumption on extinction learning using a ***S***ingle ***E***pisodic ***E***thanol (SEE) exposure model. SEE was administered 24 hours after the third conditioning trial and 24 hours before extinction training. Contextual fear conditioning trials proceeded as outlined in the methods section. Following conditioning, animals were counterbalanced into two treatment groups: Group 1 (n = 35 males; n = 30 females) received ethanol (EtOH), while Group 2 (n = 33 males; n = 30 females) received saline. A two-way repeated measures ANOVA revealed no significant differences in freezing time between the groups, but there was a significant difference across trials [F (3.331, 897.4) = 138.9, p < 0.0001], indicating that both groups had comparable capacity for conditioned stimulus/unconditioned stimulus (CS/US) pairing (Fig. 2A-B).

**Figure 2:**
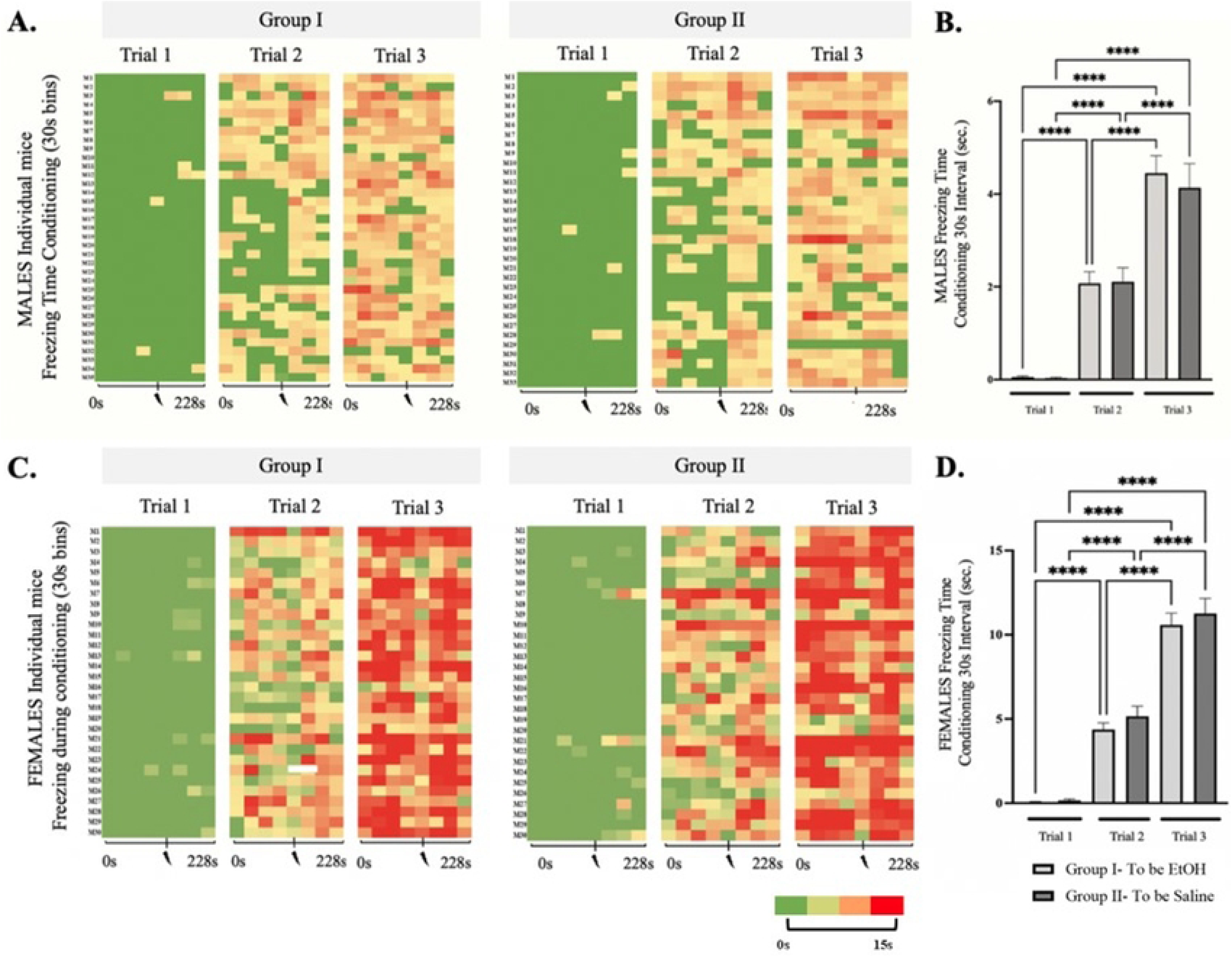
Context Fear Conditioning before SEE exposure in adolescent male and female mice. Histogram of Trial 1, Trial 2, and Trial 3 of Context Fear Conditioning of the **(A.)** male and **(C.)** female mice that will be injected with 20% EtOH (Group I: males: N=35, females: N=30) and Saline (Group II: males: N=33, females: N=30). The colors represent, in a 30 seconds’ scale, the amount of time they spent in freezing behavior. Green represents 0 second and Red represents 15 seconds or more. Mean time of Freezing for Trial 1, Trial 2, and Trial 3 of Context Fear Conditioning for **(*B)*** males and (***D)*** females in a 30s interval after conditioning prior to i.p. injections for mice that will later be segregated into saline and 6hr EtOH groups. Repeated measures two-way ANOVA, Bonferroni post hoc, **** p ≤ 0.0001. Error bars show standard error of the mean (S.E.M.).

For male mice, during the first trial of conditioning, freezing times were 0.05 ± 0.03 sec and 0.03 ± 0.02 sec; for the second trial, 2.07 ± 0.24 sec and 2.11 ± 0.30 sec; and for the third trial, 4.45 ± 0.37 sec and 4.13 ± 0.51 sec for Group 1 and Group 2, respectively, with p-values > 1.0 within 30- second bins. Final conditioning values represent approximately 15% freezing time in response to contextual fear conditioning after three trials of a single shock per trial, consistent with previously published results (33).

Similarly, in adolescent female mice, two-way repeated measures ANOVA showed no significant differences in freezing time between the groups, but a significant difference between conditioning trials [F (3.468, 728.2) = 482.6, p < 0.0001] (Fig. 2C-D). Both groups displayed comparable learning capacity, with freezing times of 0.06 ± 0.02 sec and 0.16 ± 0.07 sec (p = 0.51) for the first trial; 4.37 ± 0.39 sec and 5.16 ± 0.59 sec (p = 0.27) for the second trial; and 10.58 ± 0.69 sec and 11.26 ± 0.89 sec (p = 0.79) for the third trial, for Group 1 and Group 2, respectively, during conditioning.

Significant differences were observed between male and female mice, with females exhibiting higher freezing times per conditioning trial compared to males [F (5.167, 1306) = 285.2, p < 0.0001] (Fig. 3A). Previous studies have reported sex differences in freezing behavior, with females showing increased freezing and higher anxiety levels (55, 56), including in context-based fear conditioning (57, 58). Other studies have reported higher freezing responses in males (59–62). Differences in age groups of the mice and conditioning parameters may be responsible for the varying responses. In our study, females showed higher freezing times during the third conditioning trial, with significantly greater freezing percentages compared to male mice [F (3, 90) = 33.79, p < 0.0001] (Fig. 3B). Similar to previous studies, this has been ascribed to both the influence of specific hormonal profiles in females estrous cycle stages (58) and differential competition between hippocampus and amygdala-dependent processes in male vs. female mice (57).

**Figure 3:**
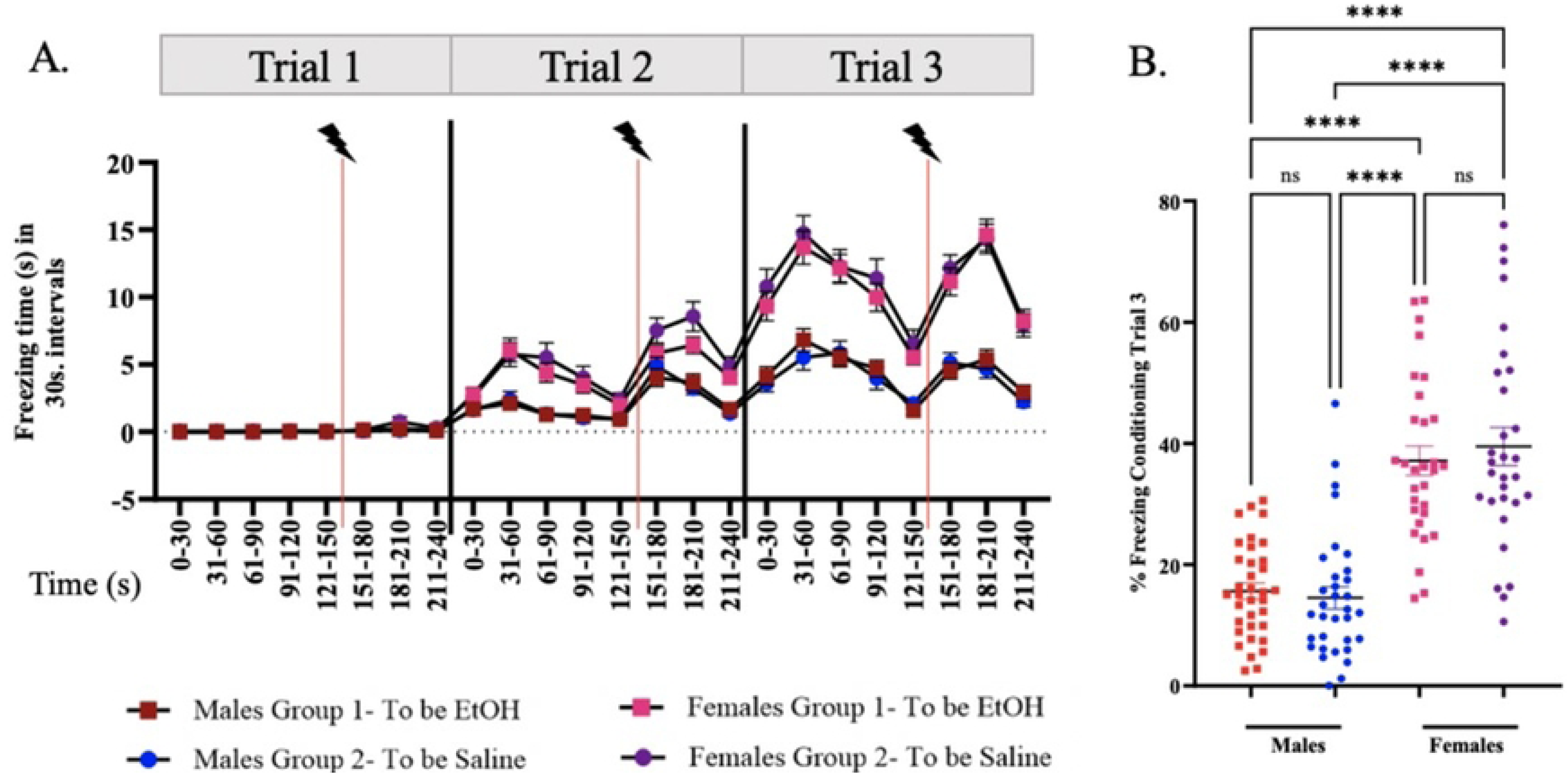
Comparison of Males vs Females during Context Fear. **(A)** Line graph comparing males’ vs females’ freezing times during the 3 conditioning trials using 30s bins. Volt with red line indicate foot shock. **(B)**. Dot Plot comparison of percent freezing during conditioning trial 3 between male and female groups. Repeated measures two-way ANOVA, Bonferroni post hoc, **** p ≤ 0.0001. Error bars show standard error of the mean (S.E.M).

To assess anxiety during conditioning, we measured the time spent in the center and periphery of the conditioning chambers (Fig. 4A-B) during CS/US pairing. Untreated male mice showed no differences between Group 1 and Group 2, nor in time spent in the center versus the periphery (Fig. 4C). In contrast, although there were no differences between Group 1 and Group 2 in untreated female mice, females spent significantly more time in the periphery compared to the center [F (1,178) = 25.77, p < 0.0001] (Fig. 4D) during context fear conditioning. This suggests that the observed differences in freezing behavior may be related to increased anxiety.

**Figure 4:**
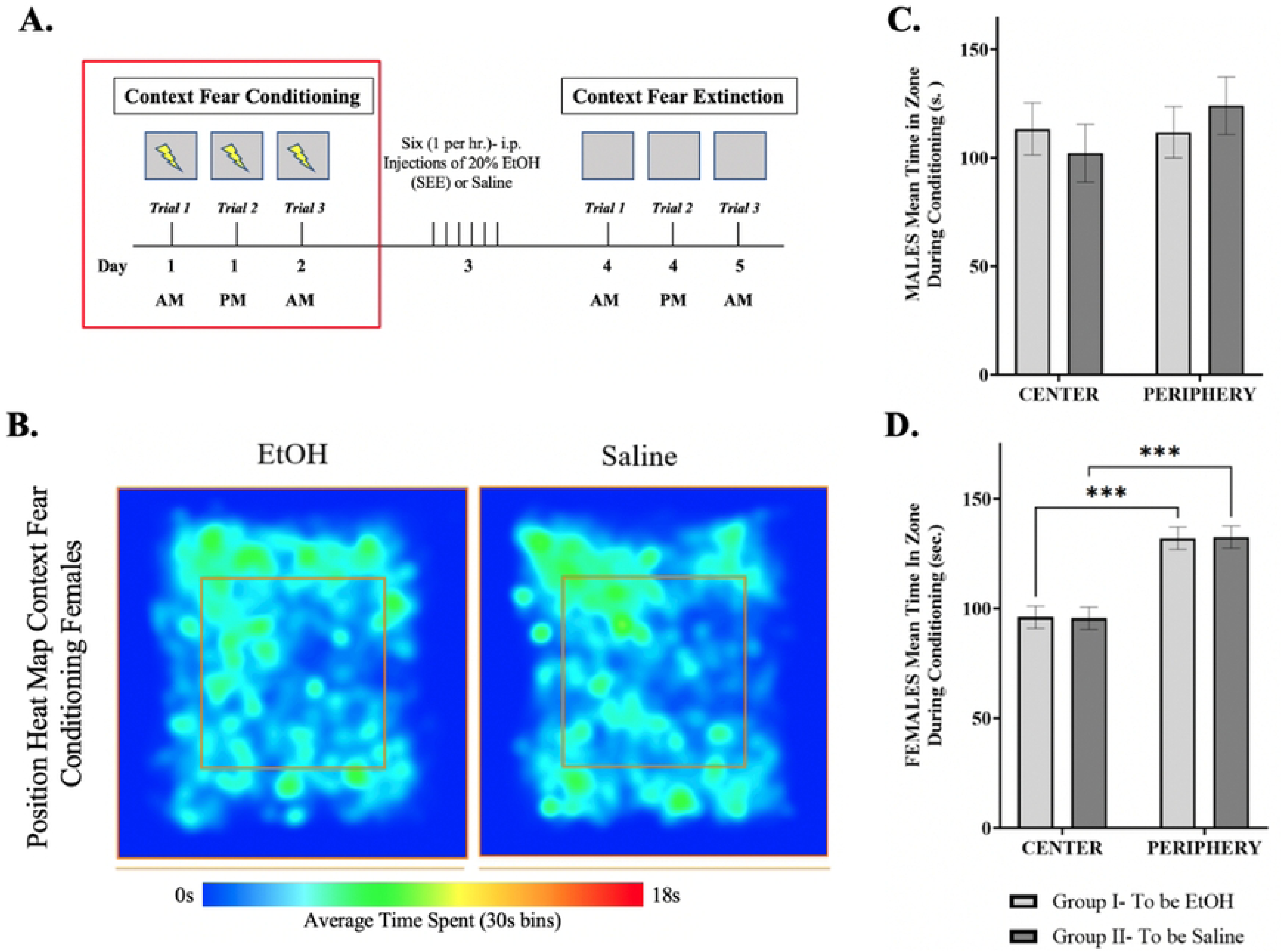
Time spent in zones during extinction trials. (***A.***) Representative diagram showing the time frame in which the data was obtained. (***B.)*** The trajectories in the heat map represents the average time spend in specific areas within the center (area within the inner box – red outline) or periphery (area surrounding the inner box) with blue representing 0s incrementally increasing until reaching red constituting 18s (maximum time in the location) using the ANY-maze video tracking analysis software system. Mean time spent in center and periphery of Context Fear Conditioning for **(*C)*** males and (***D)*** females in a 30s interval. Repeated measures two-way ANOVA, Bonferroni post hoc, *** p≤ 0.001. Error bars show standard error of the mean (S.E.M.).

### Contextual Fear Extinction Learning

Context fear extinction learning was assessed 24 hours after exposure to single episodic ethanol (SEE) exposure. In male adolescent mice, extinction learning was evident as shown by a significant decrease in freezing time between trials in the control (saline-treated) group. However, ethanol (EtOH)-treated mice exhibited consistent extinction deficits, specifically during the third extinction trial [F (4.002, 1040) = 10.011, p < 0.0001] (Fig. 5B). These findings highlight the impact of a single binge-like episode on behavioral changes in adolescent male mice.

**Figure 5:**
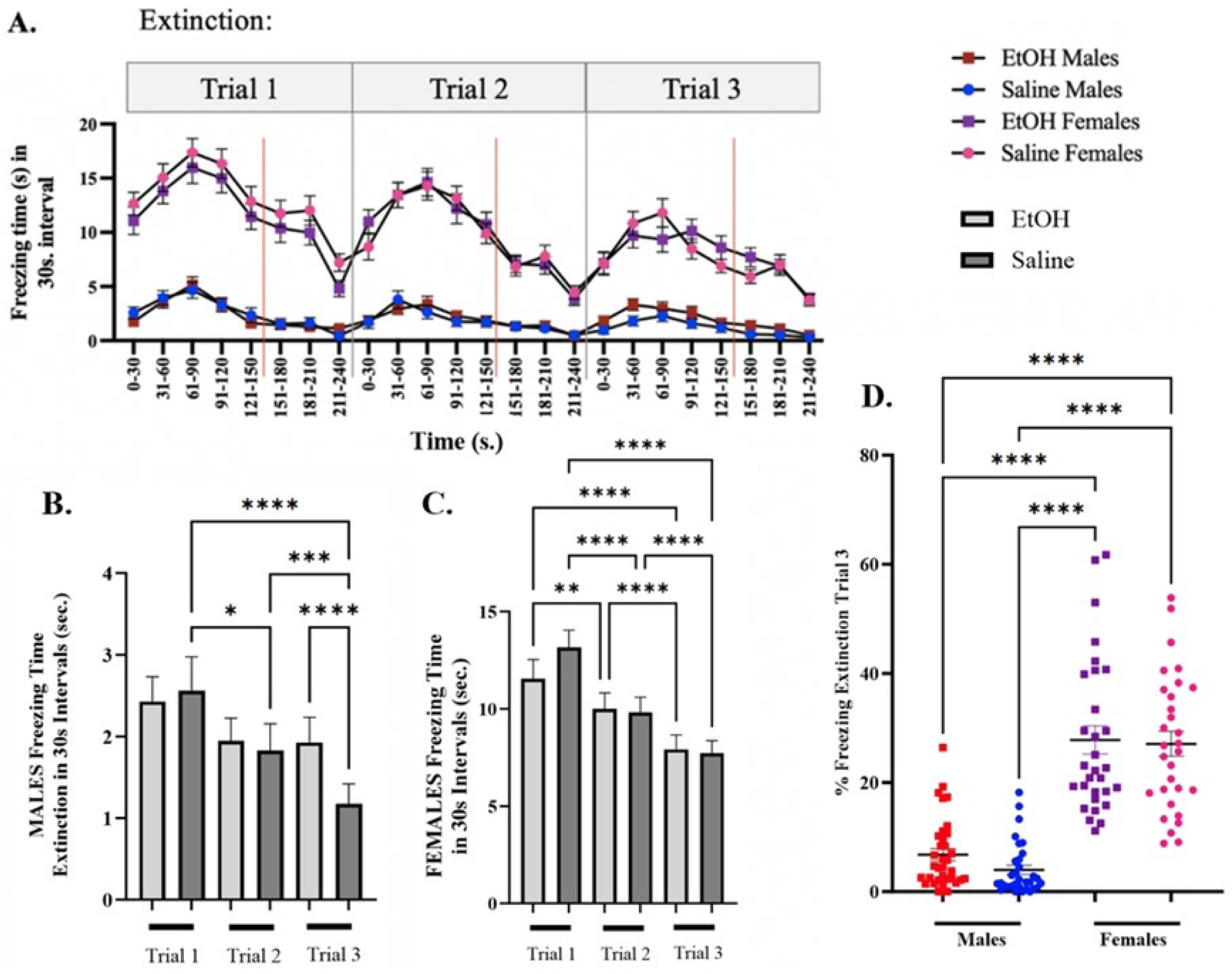
Context Fear Extinction after SEE exposure in adolescent male and female mice. **(A.)** Line graph comparing males’ vs females’ freezing times during the 3 extinction trials. Mean time of Freezing for Trial 1, Trial 2, and Trial 3 of Context Fear Extinction for **(*B.)*** males and (***C.)*** females in a 30s interval for mice that were segregated into saline and 6hr EtOH groups. **(D.)** Dot Plot comparison of percent freezing during extinction trial 3 between male and female groups. Repeated measures two-way ANOVA, Bonferroni post hoc, * p ≤ 0.05, *** p≤ 0.001, **** p ≤ 0.0001. Error bars show standard error of the mean (S.E.M.).

Interestingly, no differences in extinction were observed between treatment groups in the female adolescent cohort. Regardless of treatment, a significant decrease in freezing behavior across trials was observed [F (4.811, 1010) = 62.18, p < 0.0001] (Fig. 5C). During the first extinction trial, saline-treated females froze for 13.15 ± 0.89 seconds compared to 11.56 ± 0.90 seconds in the EtOH-treated group. In the second trial, freezing times were 9.81 ± 0.79 seconds for saline versus 9.99 ± 0.83 seconds for EtOH, and during the third trial, 7.72 ± 0.65 seconds for saline versus 7.92 ± 0.74 seconds for EtOH. Overall, female mice exhibited higher freezing times across the three extinction trials compared to males [F (6.054, 1504) = 250.3, p < 0.0001] (Fig. 5A).

Additionally, significant sex differences were observed in freezing behavior during both conditioning and extinction, with female mice exhibiting more freezing than males throughout the experiment [F (3, 89) = 52.39, p < 0.0001] (Fig. 5D). While female mice did not show deficits in extinction of fear, the pronounced differences in freezing behavior between males and females complicate the direct comparison of the effects of alcohol exposure between the sexes.

Anxiety measures during extinction were based on time spent in the center of the chamber vs the periphery (Fig. 6A-B). EtOH treated and saline male mice spent more time in the center of the chamber than the periphery [F (1,67) = 13.67, p=0.0005] (Fig. 6C). On the contrary, both EtOH treated and saline female mice spent more time in the periphery than in the center [F (1,178) = 31.92, p < 0.0001] (Fig. 6D) during context fear extinction.

**Figure 6:**
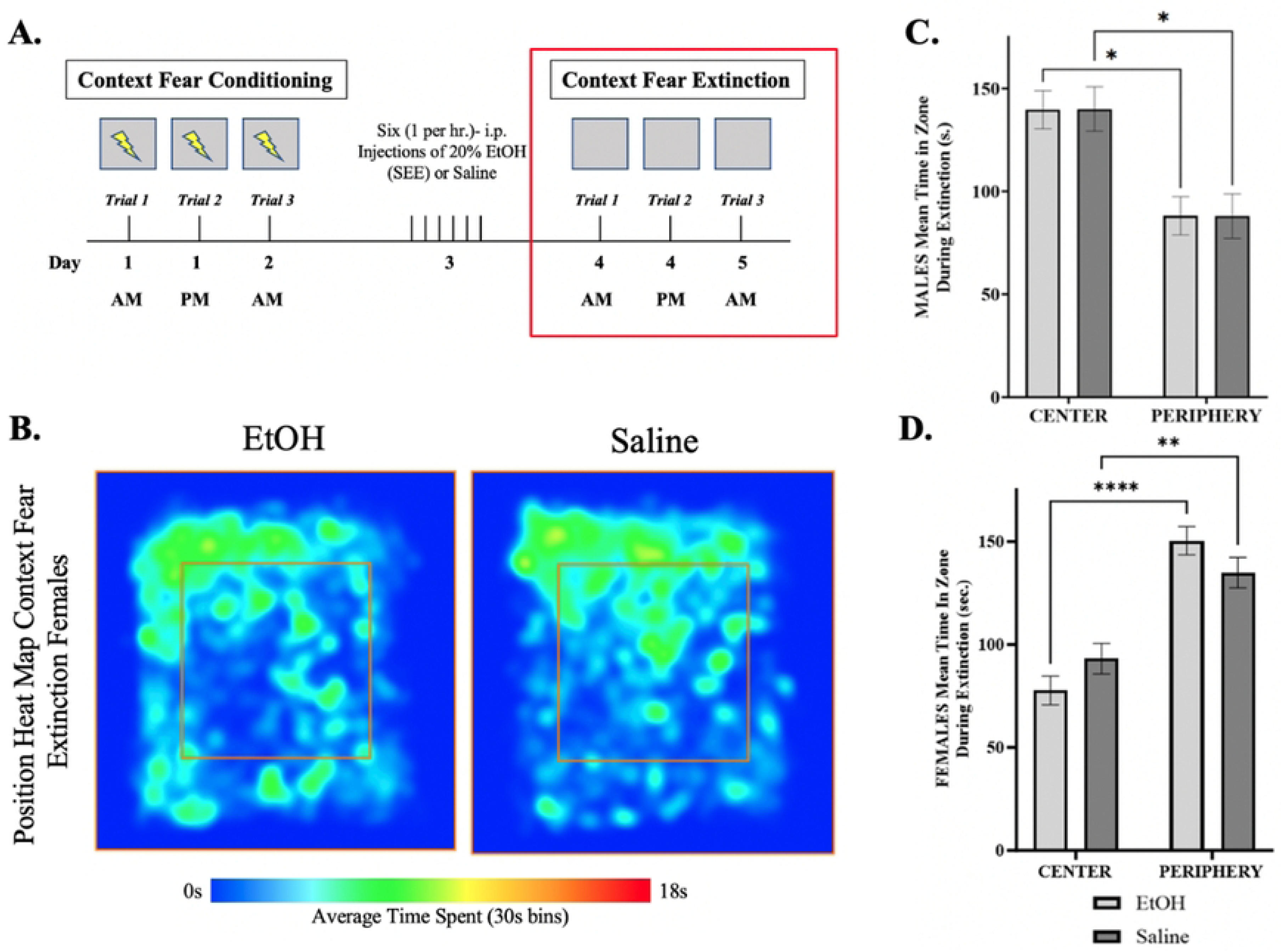
Time spent in zones during extinction trials. **(A.)** Representative diagram showing the time frame in which the data was obtained. **(B.)** The trajectories in the heat map represents the average time spend in specific areas within the center (area within the inner box – red outline) or periphery (area surrounding the inner box) with blue representing 0s incrementally increasing until reaching red constituting 18s (maximum time in the location) using the ANY-maze video tracking analysis software system. Mean time spent in center and periphery of Context Fear Extinction for **(C.)** males and **(D.)** females in a 30s interval. Repeated measures two-way ANOVA, Bonferroni post hoc, * p ≤ 0.05, **** p ≤ 0.0001. Error bars show standard error of the mean (S.E.M.).

### Open Field Test

To examine whether SEE exposure is sufficient to elicit anxiety and locomotion changes in male mice given the extinction deficits observed, we performed the open field test (OFT) (Fig. 7A) (63). Results showed no differences in locomotion behavior after SEE exposure compared to controls (Fig. 7B). However, EtOH treated mice spent significantly less time in corners: 278.0 ± 3.5 sec. versus saline controls 285.4 ± 3.2 sec. and conversely more time in the center (Fig. 7C & D). These results suggest decreased anxiety after a 24-hour withdrawal period.

**Figure 7:**
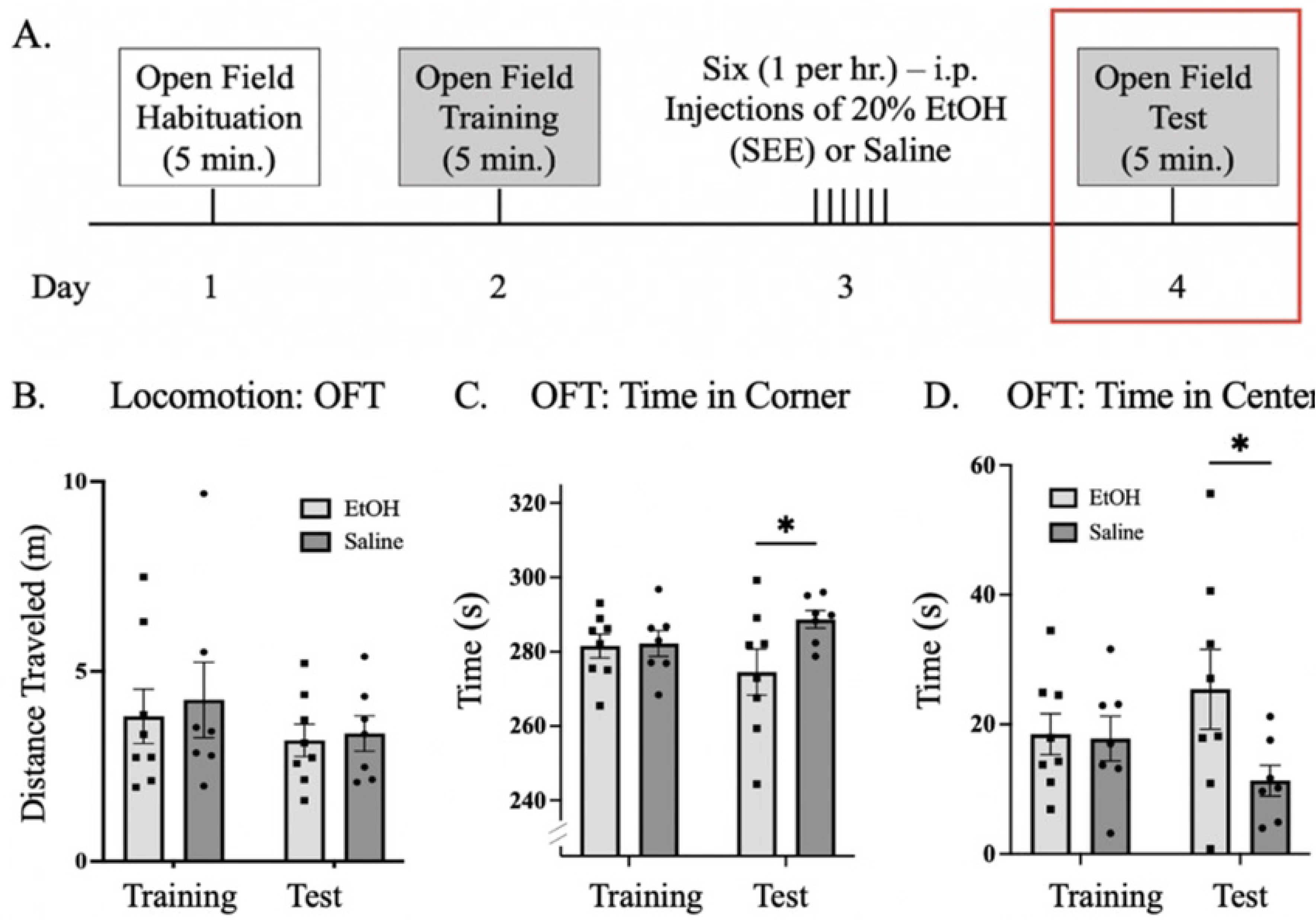
Open field test (OFT) in male mice after SEE exposure. **(A.)** Representative diagram of the timeline of the Open Field Test sessions. **(B.)** Bar graph showing distance travelled during training and test sessions. **(C.)** Bar graph showing time spent in corners in the open field arena during the training and testing sessions. **(D.)** Bar graph showing time spent in the center of the open field arena during training and testing sessions. Repeated measures two-way ANOVA, Bonferroni post hoc, * p ≤ 0.05. Error bars showing standard error of the mean (S.E.M.).

### Histology: dHPC, BLA and Striatum

After contextual fear extinction trials, we measured FKBP5 protein expression, due to its association with PTSD and anxiety-like behaviors, within the dHPC, BLA and striatum of previously injected mice. In all three regions the expression of FKBP5 protein was significantly decreased in EtOH-exposed mice (Fig.8E & 8F and Fig. 9B & 9D). This is consistent with studies linking a decrease in expression with increased risk of PTSD (64–68) in humans and fear extinction deficits in mice (47, 69, 70). We further measured the ß-catenin and GSK-3ß protein expression, due to its association with alcohol molecular tolerance (71), within the dHPC and BLA of previously injected mice (Fig.8B-D & 8G-I). The results, similarly, showed a significant decrease in expression 24hrs after extinction in both dHPC and BLA brain regions in EtOH-treated as opposed to control saline. Within the striatum, ß-catenin showed a marked tendency towards decrease but only significantly within the NAc core and shell (Fig. 9C) while GSK-3ß was significantly decreased in all four regions (Fig. 9A). This is consistent with ß-catenin expression involved in extinction learning within the NAc where studies have linked increased levels of NAc ß-catenin expression to facilitated extinction learning (72). Conversely, blocking its expression within the NAc resulted in impaired extinction.

**Figure 8:**
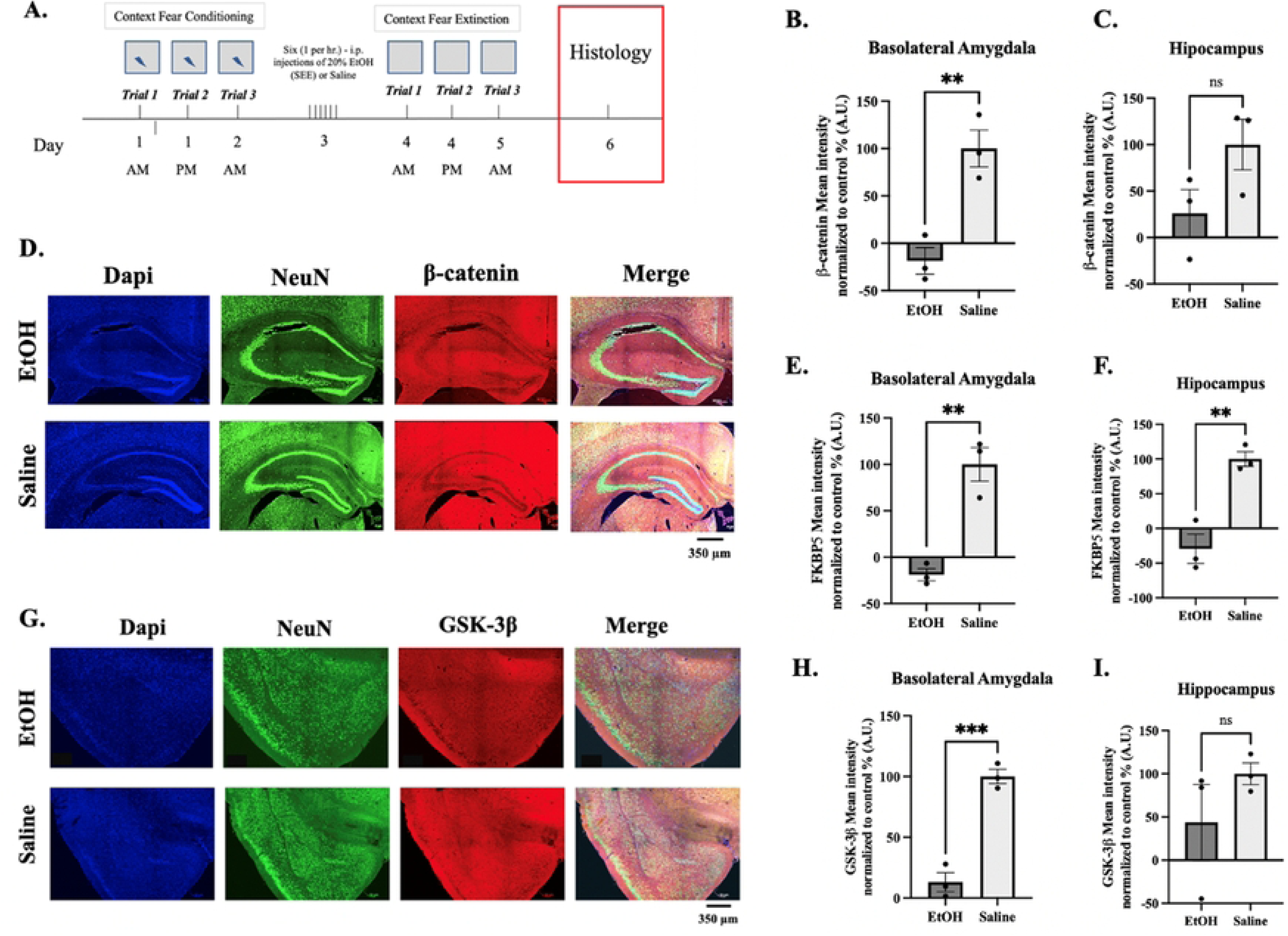
Histology of coronal sections of mouse brains after Context Fear Extinction measuring expression of FKBP5, GSK-3ß and ß-catenin. **(A)**. Representative diagram of the timeline of histology procedures. Bar graphs of quantification of the mean intensity protein expression normalized to control (%) for ß-catenin **(B.)** Basolateral amygdala (BLA), **(C.)** dorsal Hippocampus (dHPC), FKBP51 **(E.)** BLA, **(F.)** dHPC, GSK-3ß, **(H.)** BLA, and **(I.)** dHPC (N=3, S.E.M.). **(D.)** Representative confocal image of dHPC coronal slice stained with DAPI (blue – cell body), NeuN (green–neuronal marker), ß-catenin (red) and all three fluorophore images merged from mice i.p. injected with EtOH or Saline. **(G.)** Representative confocal image of BLA coronal slice stained with DAPI (blue – cell body), NeuN (green–neuronal marker), GSK-3ß (red) and all three fluorophore images merged from mice i.p. injected with EtOH or Saline. Asterisks represent statistically significant difference “ns” p > 0.5, * p ≤ 0.05, and *** p ≤ 0.001. Error bars showing standard error of the mean (S.E.M.). Measure bar represents 350 µm.

**Figure 9.**
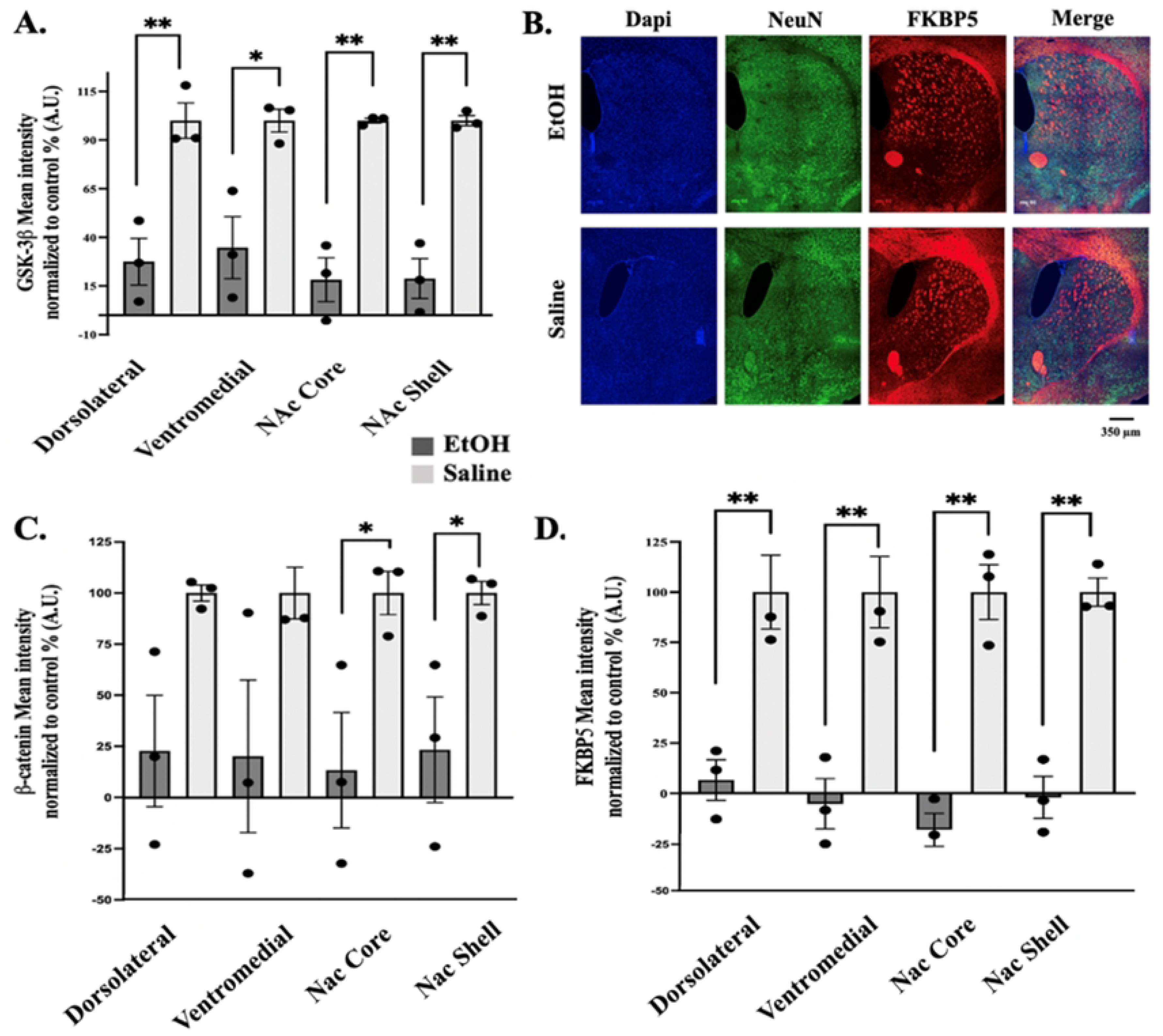
Protein expression within striatal coronal slices after Context Fear Extinction. **(A.)** Bar graph showing the mean intensity percentage normalized to control in brain regions (EtOH N=4 and Saline N=3) within the striatum; dorsolateral striatum, ventromedial striatum, NAc Core and NAc Shell. **(B.)** Representative images of the striatum in EtOH and Saline injected mice after context fear extinction. In blue is mounting media with DAPI (cell body), in green is NeuN (neuronal marker) and in red is FKBP51 (protein expression). **(C.)** Bar graph quantifying protein expression as a mean intensity normalized to control values (%) of β-catenin (EtOH N=3 and Saline N=3), **(D.)** FKBP5 (EtOH N=3 and Saline N=3). Asterisks represent statistically significant difference “ns” p > 0.5, * p ≤ 0.05, and *** p ≤ 0.001. Error bars showing standard error of the mean (S.E.M). Measure bar represents 350 µm.

### Every-Other- Day Drinking

Experiments designed with EOD prior to conditioned fear and extinction trials (Fig. 10A) showed SEE-treated male mice consumed less and preferred less EtOH than their Saline-injected controls. EtOH consumption prior to extinction was 31.76 ± 1.49 g/kg for mice treated with SEE exposure and 36.27 ± 1.74 g/kg for Saline controls, with a statistical significance of [F (2,26) =3.76, p=0.37] between treatments (Fig. 10B). EtOH preference was higher for Saline-treated mice with [F (1,28) =12.68, p= 0.0013] (Fig. 10D), while water consumption was no different between treatment groups (Fig. 10C).

**Figure 10:**
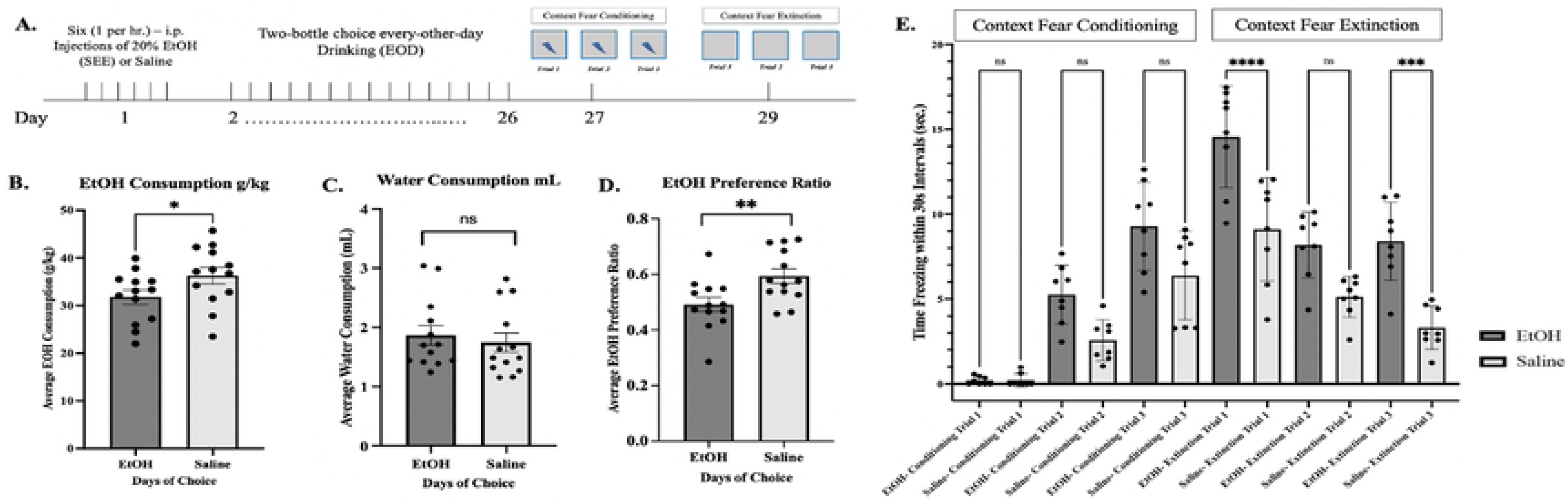
SEE and EOD prior to Context Fear Conditioning and Extinction. **(*A*.)** Schematic depicting timeframe for experimental design. **(*B*.)** Average EtOH consumption per day in g/kg for mice treated with EtOH SEE protocol and Saline control, N=12 and N=13 respectively. **(*C.)*** Average water consumption (73) for EtOH and Saline. **(*D*.)** Average EtOH preference ratio for SEE treated mice and Saline. **(*E.)*** Time freezing within 30 sec. intervals with a maximum freezing of 31% for SEE treated animals during conditioning. Extinction of SEE treated mice plateau at 28%. Maximum conditioned freezing of Saline controls 21% with a maximum extinction of 11%. Repeated measures two-way ANOVA, Bonferroni post hoc, “ns” p > 0.5, * p ≤ 0.05, *** p≤ 0.001, **** p ≤ 0.0001. Error bars showing standard error of the mean (S.E.M.).

### Contextual Fear Conditioning and Extinction after EOD

In our study alcohol pre-exposure had no significant effects on contextual fear conditioning (Fig. 10E). However, extinction learning was, once again, significantly decreased in SEE-treated male mice during extinction trials 1 & 3. Trial 1 results show 14.56 ± 1.06 seconds of freezing time for EtOH treated mice and 9.12 ± 1.08 seconds for Saline controls quantified in 30 second bins. Trial 3 results showed EtOH mice freezing 8.41 ± 0.81 seconds and 3.31 ± 0.45 seconds for Saline. ANOVA Two-Way statistical analysis showing, [F (11,84) = 34.38, p=0.0015] difference between treatments. Conditioned learning was characteristically incremental for both treatments. During extinction Saline-treated mice decreased in freezing time incrementally, as well. However, with EtOH-treated mice, freezing time during extinction plateaued during the second and third extinction trials suggesting a clear deficit in extinction learning due to SEE pre-exposure.

### Extinction Period in Novel Context

To observe the relevance of context and ethanol exposure on fear memories, adolescent male mice were exposed to different context while in conditioning and extinction (Fig. 11A). Ethanol exposed male mice demonstrated significant differences after conditioning trial 3 in Context A and extinction trials 1-3 in Context B (Fig. 11B). These results suggest associative learning was specific to the original context, context A, given that the switch to context B did not elicit recall of the original fear response. Importantly, extinction is strongly context specific, while conditioning is typically less so (74); therefore, deficits in extinction observed 24 hours after SEE treatment (Fig. 5B) are likely specific and hippocampus dependent.

**Figure 11:**
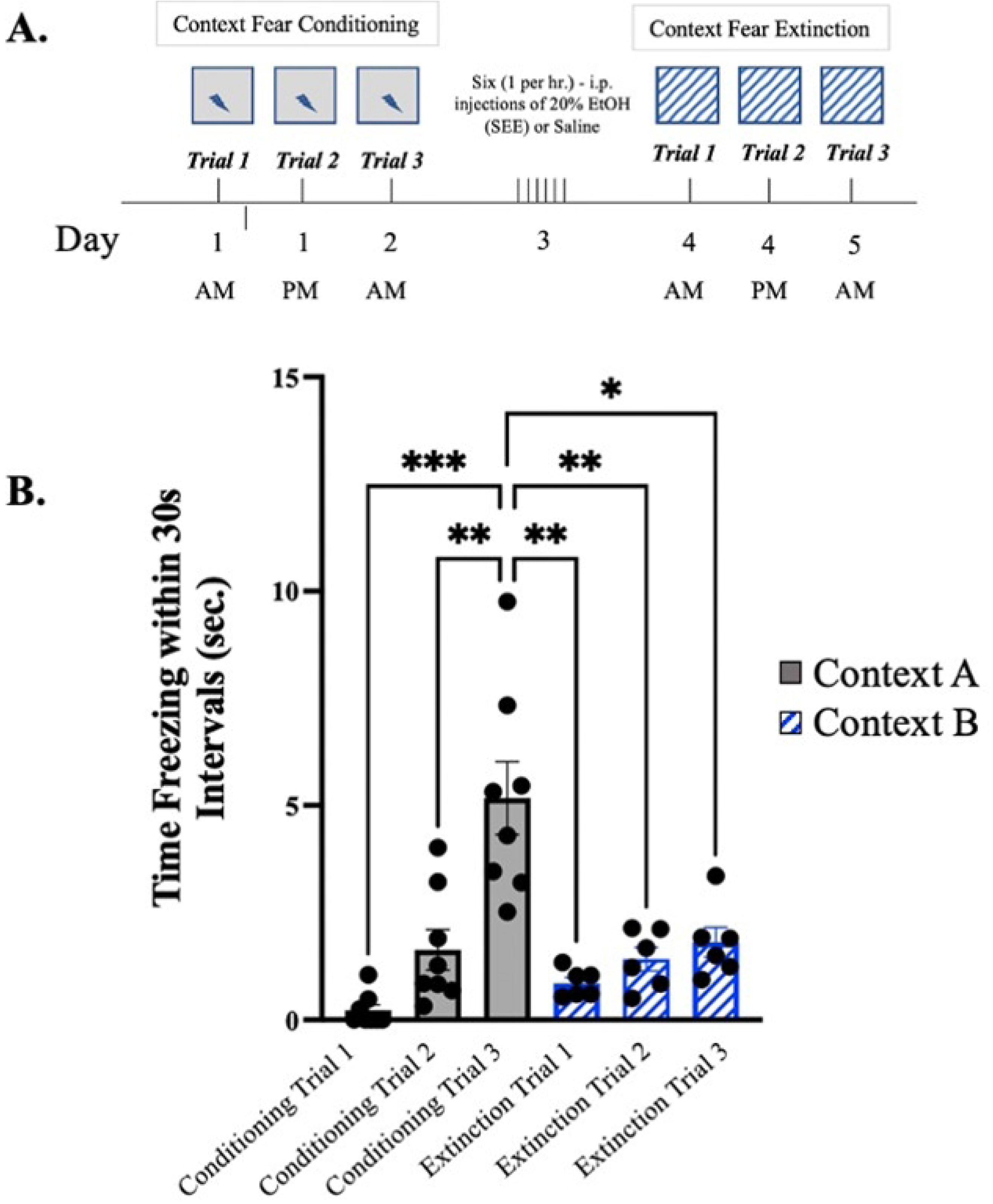
Context Fear Extinction trials under different context. **(*A.)*** Schematic depiction of experimental design where grey box represents “context A” under which original experiments took place contrasted by patterned box representing “context B” to determine context specific association. **(*B).*** Mean freezing time in 30s intervals of mice undergoing conditioning in “context A” and extinction in “context B”. Repeated measures two-way ANOVA, Bonferroni post hoc, “ns” p > 0.5, * p ≤ 0.05, *** p≤ 0.001, **** p ≤ 0.0001. Error bars show standard error of the mean (S.E.M.).

## DISCUSSION

Individuals with post-traumatic stress disorder (PTSD) experience significant psychiatric distress, characterized by the re-experiencing of a traumatic event even when the threat is no longer present. Approximately 44.7 million people will encounter PTSD at some point in their lives, with around 40% of these cases also involving co-occurring alcohol misuse issues. Despite the strong correlation between PTSD and alcohol misuse, the interplay between these two conditions remains poorly understood. Studies have shown that prior exposure to traumatic events can predict an increase in both PTSD symptoms and binge drinking among adolescents (21, 24, 75, 76). PTSD is often studied by assessing deficits in the extinction of fear responses following classical fear conditioning, such as when a tone or contextual stimulus no longer predicts a foot shock during an extinction session. Research indicates that administering high doses of ethanol before an extinction session or after several days of chronic alcohol exposure impairs extinction learning (13, 25, 29, 32, 65, 77–79). A significant barrier to advancing prevention and treatment strategies for PTSD is the absence of appropriate behavioral models that reflect clinically relevant drinking patterns. Specifically, little is known about the impact of the most common drinking behavior among young adults—single episodic or binge-like drinking—after experiencing a traumatic event and using moderate concentrations of alcohol. This study aims to explore how these prevalent drinking patterns in both male and female adolescents and young adults contribute to the development of neuropathology associated with PTSD.

### Binge-like Alcohol-induced “SEE”-induced Deficits in Fear Extinction

The timing and concentration of alcohol have distinct effects on learning and memory (29, 78, 79). Chronic alcohol exposure influences context conditioning and extinction in unique ways, potentially leading to more significant and lasting alterations in fear extinction processes, even after repeated exposure to contexts where fear has been extinguished (25, 77). Extended alcohol consumption can modify neural circuits associated with fear extinction, particularly in the prefrontal cortex and amygdala, contributing to maladaptive fear responses and an increased susceptibility to anxiety disorders (25, 65, 80). Additionally, acute alcohol exposure before context extinction trials has been found to cause deficits in extinction learning. These deficits may arise from ethanol’s interference with the formation of extinction memories or from the association of ethanol’s aversive properties with the context itself (32, 78). Understanding these differences is essential for developing effective interventions for individuals facing alcohol use disorders and co-occurring anxiety conditions.

To model a binge-like episode commonly associated with the co-occurrence of PTSD and AUD (81), we selected a single episodic exposure lasting 6 hours, administered 24 hours before extinction trials. This *in-vivo* exposure, binge-like alcohol exposure (SEE), elicits persistent neuronal adaptations known as alcohol molecular tolerance involving changes in BK channel surface expression in mice (50, 71). In mice, contextual fear conditioning increases intrinsic excitability within the dHPC (33–35), which classically relies on changes in BK-channel surface expression (82–84). The reversal of excitability increase due to conditioned learning needs to be reversed for extinction learning to proceed (33). Therefore, we hypothesized that SEE exposure persistently reinforces the increase in intrinsic excitability triggered by conditioning, disrupting its ability for reversal and subsequent extinction learning. Our results support the identification of clear extinction deficits after SEE exposure in male mice but not in female mice. The timing of the extinction impairment (within-session) suggests an alteration of processes within the BLA (85–87), rather than prefrontal cortex. Furthermore, extinction impairment could indicate that the regular function of the hippocampus in the control and regulation of the learning and memory processes is being affected by the ethanol exposure before the extinction trials (33–35, 73). Results further support the possibility that the intersecting catalytic events between alcohol exposure and deficits in extinction learning may involve the regulation of proteins involved in the Wnt/ß-catenin canonical signaling pathway expression in relevant brain circuits such as dHPC, NAc and BLA (88, 89). Thus, the most common drinking pattern among adolescents and young adults, especially after a traumatic event, could represent a previously unknown risk in the development of psychiatric disorders.

### Sex differences in male vs female mice

Sex differences in fear conditioning and alcohol use are well-documented, with women being twice as likely to develop PTSD compared to men (20% vs. 8%), and this disparity extends to the co-occurrence of alcohol use disorder (AUD) (90). While the exact mechanisms behind these sex differences remain unclear, evidence suggests that intrinsic properties and hormonal variations contribute to sex-specific fear responses (56, 91–98).

Research consistently shows that female mice voluntarily consume more alcohol (99–102). Although our behavioral model did not reveal extinction deficits in females, we observed higher freezing behavior in females, which may indicate increased anxiety (Fig. 3A). This aligns with recent findings that suggest differences in fear conditioning and anxiety are linked to the differential recruitment of the dorsal hippocampus and basolateral amygdala (BLA) (103). Ethanol exposure has also been shown to increase anxiety-like behavior in females by enhancing locomotor activity in a prelimbic cortex-dependent manner in adolescent female mice (80).

In terms of neural mechanisms, the hippocampus, amygdala, and prefrontal cortex—key regions involved in fear learning and extinction—may function differently between sexes. Estrogen, for example, can modulate synaptic plasticity in these brain areas, potentially leading to prolonged fear responses in females (60, 61). Hormones play a crucial role in fear responses, especially in females, where elevated estradiol levels and apolipoprotein E4 (E4) expression have been shown to enhance previously learned fear memories, suggesting a stronger fear memory formation in females (92, 96). Moreover, E4 has been associated with alcohol-induced memory impairment, raising the possibility that ethanol exposure could exacerbate extinction learning deficits in female mice through E4-related mechanisms (96, 104, 105).

While studies have demonstrated sex differences in the effects of ethanol on fear conditioning and extinction (77, 78, 106, 107), many questions remain unanswered. Age-dependent changes in response to ethanol also appear significant (79, 80). Future research will focus on the influence of female hormone cycling on these behavioral paradigms, aiming to clarify how different hormonal phases impact extinction learning in the context of alcohol exposure.

### Role of FKBP5, GSK-3β, and ß-catenin in Behavioral Adaptations

A critical aspect of studying comorbidity is understanding how protein expression changes in response to both alcohol use disorder (AUD) and stress or trauma. FKBP5 is a gene encoding a molecular co-chaperone of the glucocorticoid receptor (GR), which modulates intracellular glucocorticoid signaling and plays a vital role in regulating the stress response (37–41). Co-chaperones of the steroid receptor complex alter GR sensitivity, leading to a prolonged stress response (37, 38, 40), and increasing the risk for PTSD (41, 43) as well as impacting fear memory regulation (39, 45). Moreover, modulating FKBP5 expression has been identified as a potential target for preventing stress-related alcohol consumption (42).

FKBP5 also interacts with the AKT signaling pathway, which is upstream of Glycogen synthase kinase 3 beta (GSK-3β) (69). AKT activation leads to the phosphorylation of GSK-3β, inhibiting its function and thereby promoting the activation of the Wnt/β-catenin signaling pathway. The Wnt/β-catenin pathway is critically important, especially in the context of binge-like alcohol exposure via SEE exposure, as it triggers the development of alcohol molecular tolerance (50, 71). Transient activation of β-catenin can serve as a catalyst for other key neuroplasticity events in brain regions such as the amygdala (108), striatum (17, 109), and hippocampus (110, 111).

Although we typically expect an inverse relationship between GSK-3β and β-catenin expression during Wnt/β-catenin activation, various factors can influence the final expression pattern observed post-extinction. Nonetheless, evidence points to dysregulation of proteins associated with the canonical Wnt/β-catenin pathway, likely contributing to extinction deficits induced by binge-like SEE exposure. While these changes are promising, it’s important to recognize that these proteins participate in dynamic processes, and their expression likely fluctuates during extinction learning. Given the controlled conditions across experimental groups, our findings strongly suggest that binge-like ethanol exposure significantly impacts extinction learning.

In a social defeat model of depression, reduced levels of specific forms of GSK-3β within the nucleus accumbens (NAc) were linked to increased susceptibility to chronic stress, in connection with the canonical Wnt/β-catenin signaling pathway (54, 112, 113). Activation of the Wnt/β- catenin pathway is associated with resilience to depression, while its downregulation is correlated with heightened stress responses. Our findings showed significant decreases in both GSK-3β and β-catenin expression, aligning with these observations. Transient changes in this pathway could have substantial effects throughout the experimental timeline and may contribute to the initial development of persistent alcohol molecular tolerance followed by the emergence of a stress-like phenotype within the contextual fear extinction paradigm. We hypothesize that Wnt/β-catenin pathway activation underlies the initial deficits in extinction learning, while subsequent recovery of β-catenin expression may represent a compensatory mechanism driving long-term changes.

The observed decrease in FKBP5 expression in the dorsal hippocampus (dHPC), striatum, and basolateral amygdala (BLA), associated with deficits in fear extinction, is consistent with changes seen in postmortem brain samples from PTSD patients. Additionally, effective PTSD treatment through cognitive behavioral therapy (CBT) has been linked to increased FKBP5 expression (64, 65, 67, 68, 114, 115). In mice, increasing FKBP5 mRNA levels in the amygdala improves extinction deficits, further emphasizing that reduced FKBP5 expression contributes to an enhanced stress response (47, 69, 70). Conversely, longitudinal studies of early developmental stress and alcohol exposure have shown elevated FKBP5 gene expression (46), suggesting a potential relationship between early-life exposure to stress or alcohol and later epigenetic changes. In these conditions, brain regions with high FKBP5 expression, such as the hippocampus, amygdala, dorsal raphe, and locus coeruleus, are implicated in both stress response and alcohol dependence (39, 41). Pharmacological studies have also demonstrated that the selective FKBP5 inhibitor, SAFit2, reduced alcohol consumption in stressed male rodents without affecting irritability (42). Overall, our findings align with the existing literature and suggest that FKBP5 may play a role in ethanol’s impact on fear extinction.

### Circuitry Mediating Fear Response

The acquisition, consolidation, and expression of contextual extinction memories are thought to be mediated by highly specific neuronal circuits embedded in large-scale brain networks principally located in the amygdala and hippocampus. The amygdala complex plays a major role in how alcohol and stress influence cellular physiology to produce disordered behavior while the hippocampus participates in how alcohol and stress influence cognitive function, emotional regulation, and memory formation (116–119). Furthermore, studies have shown the distinct roles of the dHPC and vHPC in fear extinction, such that dHPC inactivation accelerated extinction acquisition, while vHPC inactivation reduced fear expression during extinction and impaired extinction recall (120). Enhancing the connections and/or intrinsic excitability of the dorsal hippocampus could potentially explain the deficits in contextual extinction, as suppressing this area aids in the process of extinction learning (33, 120). Acute and repetitive alcohol exposure diminishes the time and effectiveness of fear extinction mainly by affecting the activation of the basolateral amygdala (BLA) following fear acquisition and suppressing activity in the central amygdala (CeA) suggesting that fear memories established while under the influence of alcohol might exhibit stronger associations with context (29). Thus, while certain components require the reversal of conditioning processes, other components contribute to a unique new learning specific to extinction. The stress response during extinction is further modulated by other brain regions, including the NAc, which is specifically linked to the AUD and PTSD association. Our results showed that the effects of binge-like SEE exposure are part of an extinction-specific neuronal ensemble (121) as observed when SEE is administered prior to fear conditioning. Consistent with the idea that discrete sub-networks of projection neurons interconnect brain areas involved in extinction, it has been found that inputs from the mPFC and the hippocampus differentially target ‘extinction neurons’ and ‘fear neurons’, respectively (122). Furthermore, dysregulation of dopamine (DA) release specifically from the medial ventral tegmental area (VTA) and/or inhibition of dopamine receptors in the NAc impairs extinction learning (123–126). Finally, the BLA has been carefully characterized both for fear conditioning and extinction learning and further linked to Wnt/ß-catenin signaling (89, 127). These data suggest that a dynamic modulation of Wnt/β-catenin signaling during consolidation is critical for the structural basis of long-term memory formation during acquisition and extinction of fear memory.

Our results support the hypothesis that a single episodic binge-like ethanol exposure, resulting in moderate intoxication lasting 6 hours, may increase the risk of extinction deficits associated with the development of trauma and stressor-related disorders. Thus, binge-drinking behavior observed in adolescents and young adults, particularly following a traumatic event, could represent an unexpected risk factor for the onset of psychiatric disorders. Our results further support the possibility that the intersecting catalytic events between alcohol exposure and deficits in extinction learning may involve the regulation of the WNT canonical signaling pathway and FKBP5 expression in relevant brain circuits such as the hippocampus, NAc, and BLA. This suggests a potential avenue for intervention whereby the risk could potentially be addressed through behavioral interventions, pharmacotherapeutics, or a combination thereof, interacting with newly identified targets elucidated in this study, thereby paving the way for innovative preventive and treatment strategies.

## ACKNOWLEDGMENTS

The authors thank Yma Escalona for training workshops in parafilm embedding tissue for histology. The authors also thank Drs. Christian E. Bravo Rivera, Dr. Demetrio Sierra Mercado, and Dr. Juan C. Jorge Rivera for feedback on the manuscript in progress.

